# Fear contagion in zebrafish: a behaviour affected by familiarity

**DOI:** 10.1101/521187

**Authors:** Priscila Fernandes Silva, Carlos Garcia de Leaniz, Ana Carolina Luchiari

**Affiliations:** Centre for Sustainable Aquatic Research (CSAR), Department of Biosciences, Swansea University, UK; Departament of Physiology, Universidade Federal do Rio Grande do Norte, Brazil

**Keywords:** emotional contagion, empathy, alarm response, animal model

## Abstract

Emotional contagion has recently been described in fish but whether it is affected by familiarity is not known. We tested whether the sight of a distressed conspecific elicited fear in zebrafish, and whether this was modulated by familiarity. Groups of six zebrafish were housed together in the same tanks for 7 days to create familiar conditions. The behaviour of individual fish was then recorded in paired tanks within sight of either a familiar or an unfamiliar individual, before and after distilled water or an alarm substance was added to the demonstrator, but not to the observer. As expected, addition of distilled water did not elicit any behavioural change in either the demonstrator or the observer. However, addition of an alarm cue triggered anti-predatory behaviours in the demonstrator which caused the expression of anti-predatory behaviours in the observer, suggesting the existence of fear contagion. Furthermore, the extent of fear contagion was affected by familiarity, and observers were more active, swam closer to the bottom and further away from the demonstrator when they watched a distressed familiar neighbour than when they watched an unfamiliar fish. Our results have implications for fish welfare because they show that fish can become stressed by simply watching others become stressed. They also have implications for experimental design because fish housed in separate tanks cannot be assumed to be statistically independent if they can eavesdrop on their neighbours.

## Introduction

Emotional contagion can be defined as the instantaneous matching of emotional state between an observer and a demonstrator (Nakahashi & Ohtsuki, 2018). This phenomenon has been explained through the perception-action mechanism, which postulates that the perception of a demonstrator’s state triggers a neural, unconscious and automatic representation of the same state in the observer, causing an equivalent expression of behaviours (Preston & de Waal, 2002). Emotional contagion is considered a component and evolutionary precursor of empathy (Preston & de Waal, 2002) and has been demonstrated in humans, birds and mammals (Gonzalez-Liencres et al., 2014; Reimert et al., 2014).

To evaluate emotional contagion studies have typically focussed on negative emotional states such as stress, pain and fear (Carnevali et al., 2017). A common measure of fear in rodents is freezing behaviour, which can be triggered by a mild electric shock (Lezak et al., 2017; Pisansky et al., 2017). Fear elicited in this way propagates from frightened demonstrators to naïve observers, resulting in increased frequency of freezing (Jeon et al., 2010; Knapp et. al, 2007; Knapska et al., 2006) and activation of the amygdala in the observer (Knapska et al., 2006). Moreover, it seems that how an emotion is shared between individuals is modulated by contextual aspects such as kinship, familiarity and social closeness (Liévin-Bazin et al., 2018; Preston & de Waal, 2002). For instance, when mice are paired with distressed demonstrators they tend to freeze if they had been reared together, but become more active if they come from different cages (Gonzalez-Liencres et al., 2014). In addition, pain perception in mice is more intense when observers are familiar with demonstrators than when they are strangers (Langford et al., 2006). Observers typically respond differently to signals sent by familiar and unfamiliar conspecifics (Gonzalez-Liencres et al., 2014; Jeon et al., 2010), a strategy thought to be adaptive as it can help avoid sensory overload (Hutchinson, 2005) and focuses attention on signals emitted by those neighbours that matter the most, including ‘nasty neighbours’ and ‘dear enemies’ (Müller & Manser, 2007).

A recent study has provided evidence for fear contagion in zebrafish (Oliveira et al., 2017), which suggests that this phenomenon may be conserved among social vertebrates. However, to what extent fear contagion in fishes can be affected by the degree of familiarity between demonstrators and observers is not known. Familiarity can broadly be defined as the ability to discriminate between individuals based on previous interactions, and is influenced by the time of interaction and the size of the group among fishes (Griffiths, 2003).

Here we used dyadic behavioural tests to assess if fear contagion was affected by familiarity in zebrafish. To this end, demonstrators were exposed to either distilled water or an alarm substance, known to cause a fear response on this species (Kalueff et al., 2013). Zebrafish are highly social (Gerlai, 2010), can discriminate familiar from unfamiliar fish after only 20 min of interaction (Hinz et al., 2013; Madeira & Oliveira, 2017), and form cohesive groups when under threat (Speedie & Gerlai, 2008). Therefore, we hypothesized that observers would show a heightened fear response to the sight of distressed demonstrators when they had been reared together (i.e. were familiar), than when they had been reared apart (i.e. were unfamiliar).

## Methods & Materials

### Experimental fish and husbandry conditions

Two month old, laboratory-reared zebrafish (*Danio rerio*) of homogeneous size were sourced from a local supplier and kept in four 50 L tanks (density = 2 fish/L) connected to a recirculation system for four months before testing. Water quality was maintained by mechanical, biological and chemical filtration, in addition to UV disinfection. Water temperature was kept at 28 ± 1°C, pH at 7.2 and ammonia and nitrite at recommended optimal levels for the species. Photoperiod was set at 12D:12L with the help of fluorescent lights (150 lumens) with the start of the light phase set at 7:00 hrs. Fish were fed commercial pellets twice a day (Nutricom Pet, 38% protein, 4% lipids) and *Artemia salina* once daily.

### Development of familiar and unfamiliar groups

Seven days prior to testing, a sample of 156 adults of both sexes was collected haphazardly from the four stock tanks, mixed, and allocated at random in groups of six to 26 × 20 L glass aquaria (40L × 20W × 25 H cm). Aquaria were filled with system water, were fitted with a sponge filter to maintain water quality, and had their bottom and sides covered with white plastic sheets to prevent visual contact with others groups. Food was offered twice a day, as above.

### Acclimation period

After seven days of being reared in groups of six, two individuals from either the same or different aquaria (i.e. familiar or unfamiliar conditions) were transferred to two 2L test aquaria (20L × 10W × 20H cm) placed side by side and left to acclimatize for 18 hours prior to testing, one fish serving as a ‘demonstrator’ and the other as an ‘observer’. To ensure that unfamiliar fish would not become ‘familiar’ during the acclimation period, a removable divider was placed between the two test aquaria, so that demonstrator and observer had no visual contact until the divider was lifted just prior to testing. Similarly, and in order to avoid a potential disruption of familiar dyads, these were acclimated without dividers, in full sight of each other. To test if this could have affected their subsequent behaviour, we tested 8 additional familiar dyads acclimatized with dividers, and compared their behaviour to familiar dyads acclimatized without dividers.

### Testing of fear contagion

Following the 18h acclimation period, the divider preventing visual contact was removed and the demonstrator and observer were simultaneously recorded (Sony DCR-SX45 Digital VCR) for 5 min (basal behaviour). A syringe connected to a small silicon tube was then used to remotely deliver 2 ml of either distilled water or an alarm substance to the demonstrator (delivery being allocated at random), and their behaviours were recorded at 10 minute intervals over an hour, as shown schematically in **Figure 1**. To obtain the alarm substance, one zebrafish from the stock tanks was euthanized by an overdose of clove oil (2 ml/L water), and c. 1 cm^2^ of skin from each flank was removed, macerated in 10 ml of distilled water and filtered. Fresh alarm substance was prepared every morning before testing.

**Figure 1.**
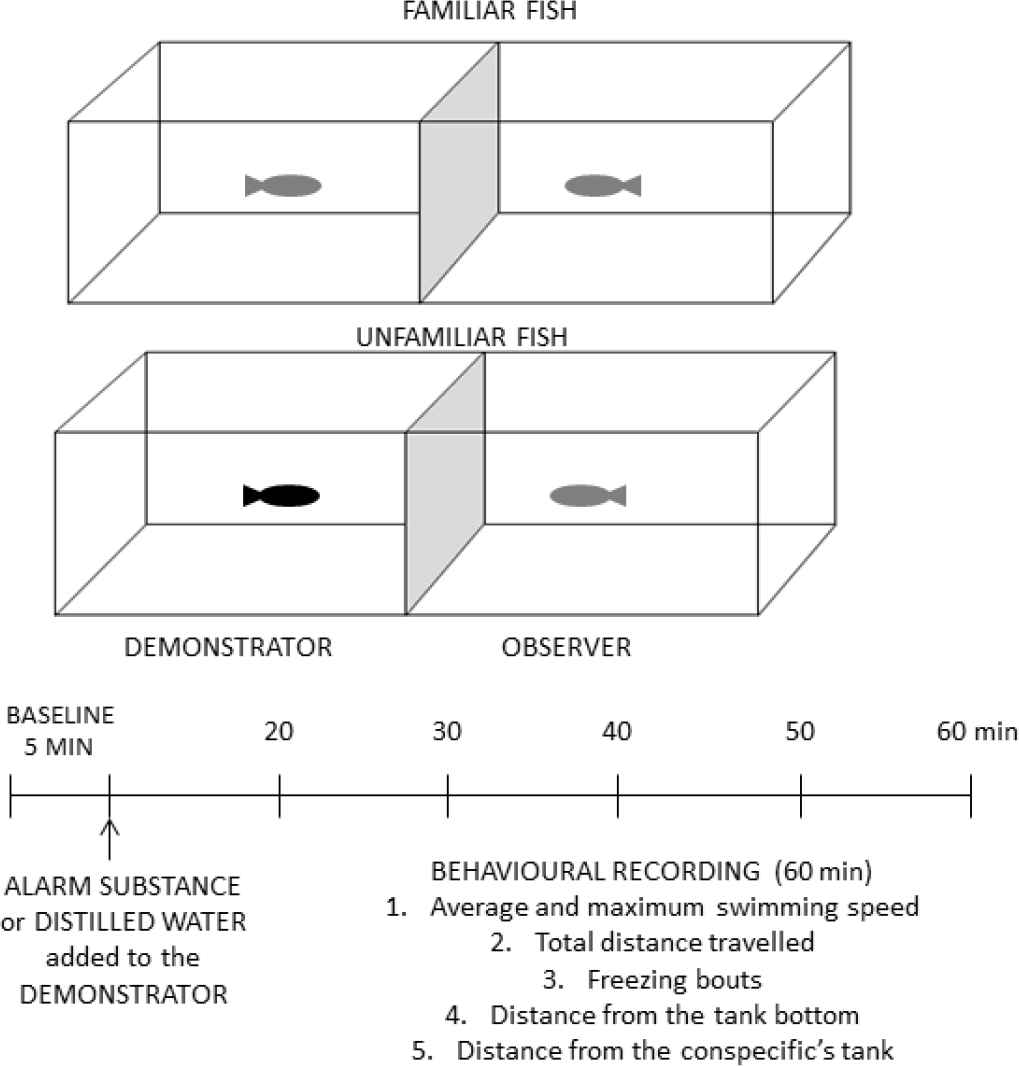
Schematic representation of experimental testing of fear contagion between familiar and unfamiliar zebrafish dyads.

We tested 48 dyads exposed to alarm substance (24 unfamiliar and 24 familiar) and 20 dyads exposed to distilled water (10 unfamiliar and 10 familiar). In addition, 20 demonstrators (10 exposed to alarm substance and 10 exposed to distilled water) were tested without observers to ascertain which behaviours were modified by the addition of the alarm substance, and to what extent the presence of observers influenced the demonstrator’s behaviour. Fish were only used once in the experiments, either as observers or demonstrators. The 10 ml of alarm substance obtained from one fish was enough to test five dyads, thus for all the experiments 12 zebrafish were used for the preparation of alarm substance.

We used *ZebTrack* (Pinheiro-da-Silva et al., 2016) to extract from the video recordings six behavioural metrics that had previously been shown to describe well the stress response of zebrafish (Gerlai et al, 2008; Luca & Gerlai, 2012; Tran & Gerlai, 2013), namely : (1) mean swimming speed, (2) maximum swimming speed, (3) total distance travelled, (4) time spent freezing, (5) swimming depth (i.e. distance from the tank bottom) and (6) mean distance to the conspecific’s tank.

### Statistical Analysis

Statistical analysis was conducted in R v. 3.4.3 (R Core Team 2013). Our experiment conformed to a fully factorial 2 × 2 × 2 BACI design (before-after-control-impact) and we modelled the behaviour of the observer (dependent variable) as a linear mixed effect model using the *lme4* (Bates et al., 2014) and *lmerTest* (Kuznetsova et al., 2017) R packages. We used as fixed effects (predictors) the behaviour of the demonstrator, the time (before or after the stressor was added), the familiarity (familiar vs unfamiliar dyad) and the stressor type (alarm substance vs distilled water), and considered the dyad identity as a random effect to control for variation among test arenas and account for potential non-independence of observations. For each behavioural metric, we started with a maximal model with all main effects and interactions and used the *step* and *dredge* functions in the *MuMIn* package (Bartoń, 2013) to arrive at a minimal adequate model via Maximum Likelihood on the basis of single deletion tests and relative changes in AICc values. The most plausible model was refitted by REML and the model adequacy and assumptions were checked by examining plots of fitted vs residuals, fitted vs observed values, as well as plots of random effects and standardized fixed effects using the *sjPlot* package (Lüdecke, 2016). We report all models within ΔAICc = 2. One dyad had missing values so there were 134 observations corresponding to 67 dyads.

To better assess the extent to which observers were able to match the overall behaviour of demonstrators, we also carried out a principal components analysis and modelled the scores along the first two first principal components (which together explained 96% of variation) as a function of fish type (observer or demonstrator) and degree of familiarity.

### Ethical note

All experimental procedures were authorized by the Animal Ethics Committee permit CEUA 042/2015 and Animal Welfare and Ethical Review Body permit IP-1516-8.

## Results

### Response to alarm cues (single tests)

Inspection of temporal data (Supplementary Material, **Figure S1**) indicated that the response of single fish to alarm cues was rapid and did not persist for more than 10 minutes after the administration of the alarm substance (probably due to habituation). We, therefore, concentrated the analysis on the first 10 minutes after addition of the stimuli. Compared to baseline values, the addition of an alarm cue resulted in an increase in average swimming speed and time spent freezing that was not observed when distilled water was added (average speed, before-after = 6.20, SE = 0.81, *t*_17_ = 7.68, *P* < 0.001; blank-alarm =7.08, SE = 1.11, *t*_27.19_ = 6.36, *P* < 0.001; after x alarm= −6.39, SE = 1.11, *t*_17_ = −5.75, *P* < 0.001; time spent freezing, before-after = 118.7, SE = 13.24, *t*_18_ = 8.97, *P* < 0.001; blank-alarm =110.6, SE = 13.52, *t*_35.94_ = 8.18, *P* < 0.001, after x alarm = −105.56, SE = 18.72, *t*_18_ = −5.64, *P* < 0.001). Fish also stayed closer to the bottom after alarm cue or distilled water were added (before-after = −3.50, SE = 1.00, *t*_18_ = −3.49, *P* = 0.002), but no significant change was detected in relation to maximum speed or distance travelled (*P* > 0.05; Supplementary Material, **Figure S1**) and these were not considered any further. Results also indicated that whether familiar demonstrators were visually isolated or not during the 18 hrs of acclimation did not affect their subsequent behaviours (Supplementary Material, **Table S1**).

### Fear Contagion from Demonstrators to Observers (dyadic tests) Swimming Speed

As expected from the single tests above, demonstrators in the dyadic tests increased their average swimming speed when an alarm substance was added, but not when distilled water was added (**Figure 2**). Observers responded by increasing their speed when the demonstrator was unfamiliar, but by decreasing it when the demonstrator was familiar (**Figure 2**; estimate demonstrator = 0.40, *P* = 0.002; estimate time = 0.93, *P* <0.001; estimate familiarity x time = 1.18, *P* <0.001; estimate time × stressor = 0.53, *P* =0.002; estimate familiarity × time × stressor = −0.60, *P* <0.001). Fear contagion, hence, was affected by familiarity.

**Figure 2.**
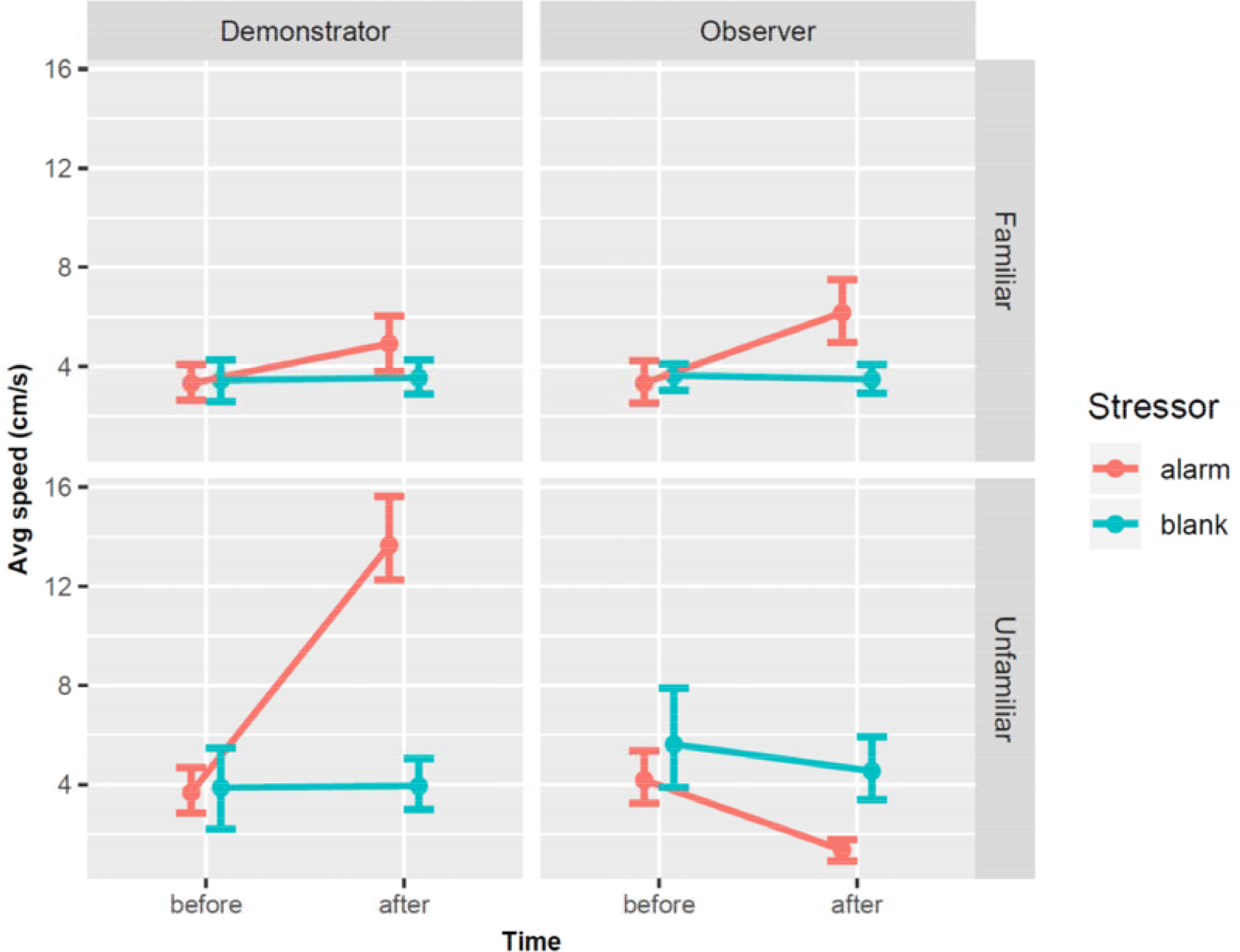
Changes in the swimming speed (mean ± 95CI) of zebrafish before (basal) and after distilled water or an alarm cue were delivered to the demonstrator in dyadic tests involving familiar and unfamiliar pairs.

### Freezing behaviour

As expected also from the single tests above, demonstrators spent more time freezing when an alarm substance was added, but not when distilled water was added (**Figure 3**). The time observers spent freezing increased when an alarm substance was added to the demonstrator (estimate = 0.85, *P* < 0.001), and also with time (estimate = 0.90, *P* < 0.001), but was negatively correlated with the time spent freezing by the demonstrators (estimate = −0.19, *P* = 0.024). There was a significant interaction between time and stressor (estimate = 0.63, *P* < 0.001) as observers only increased the time spent freezing over basal values when the alarm substance was added, not when distilled water was added. Familiarity, hence, did not influence the freezing response, which was very strong under both conditions.

**Figure 3.**
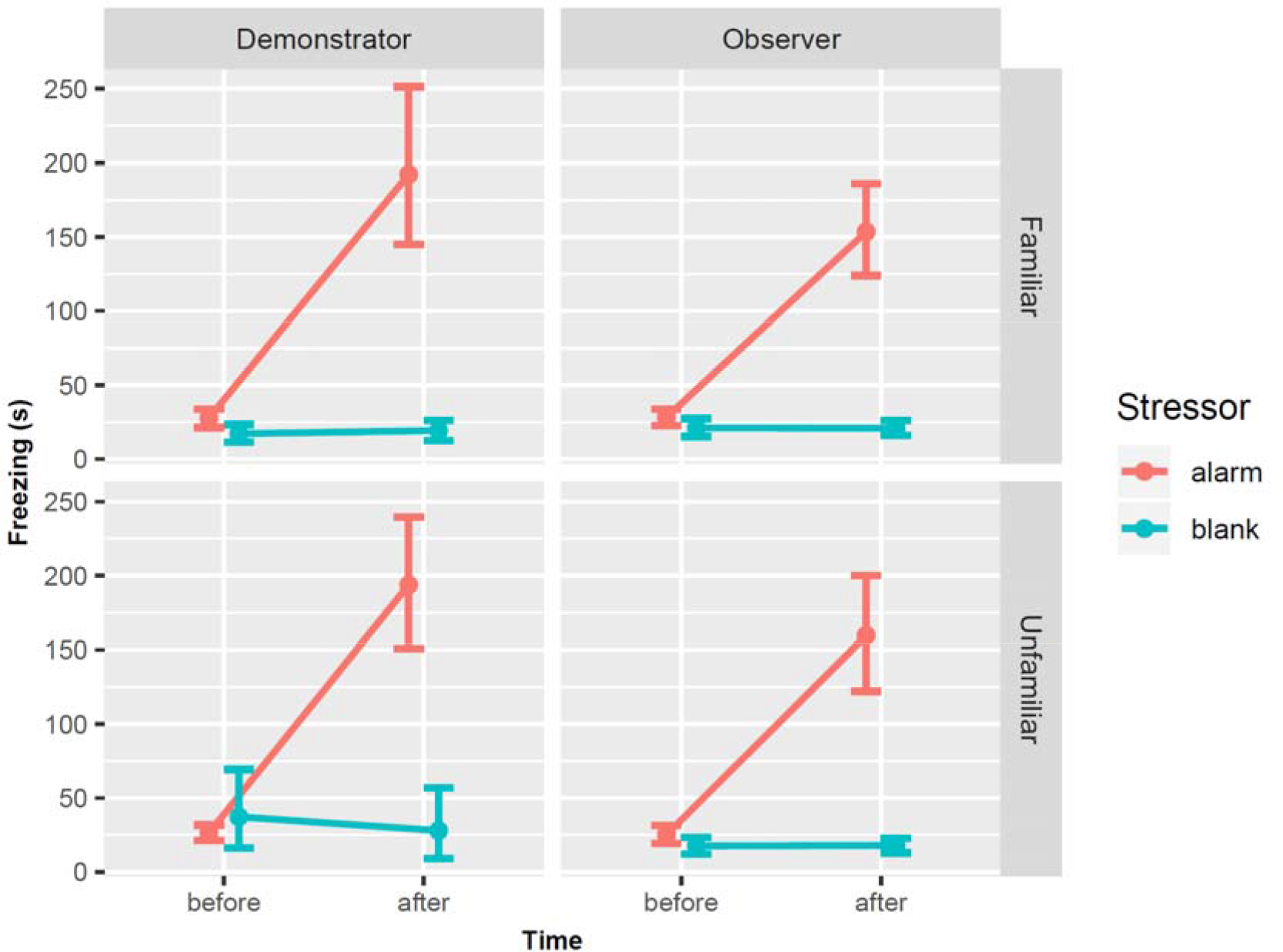
Changes in the freezing behaviour (mean ± 95CI) of zebrafish before (basal) and after distilled water or an alarm cue were delivered to the demonstrator in dyadic tests involving familiar and unfamiliar pairs.

### Distance from the tank bottom (swimming depth)

Following the addition of the alarm substance, demonstrators moved closer to the bottom of the tank, a behaviour not seen when distilled water was added. Observers mimicked this behaviour (**Figure 4**), tracking what the demonstrator did (estimate = 0.19, *P* = 0.033). Swimming depth increased over basal values (estimate = 0.23, *P* = 0.011), as well as with the addition of the alarm cue (estimate = 0.31, *P* = 0.018). There were significant interactions between demonstrator’s depth and familiarity (estimate = 0.25, *P* = 0.03), demonstrator’s depth and time (estimate = 0.25, *P* = 0.002), familiarity and time (estimate = 0.44, *P* < 0.001), familiarity and stressor (estimate = 0.54, *P* < 0.001), and familiarity × time × stressor (estimate = 0.28, *P* = 0.003). There was, hence, evidence of fear contagion which was also affected by familiarity.

**Figure 4.**
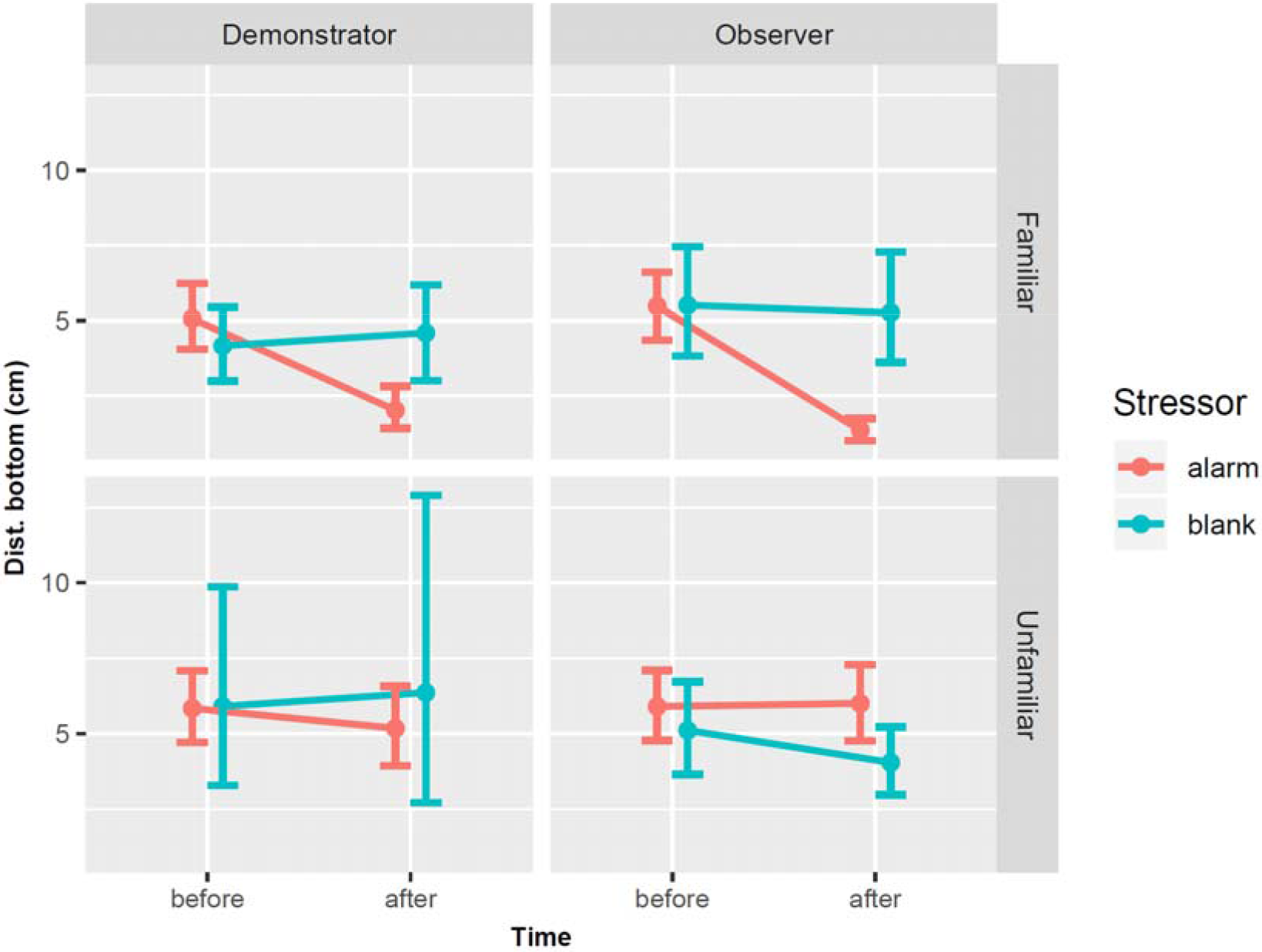
Changes in the swimming depth (mean ± 95CI) of zebrafish before (basal) and after distilled water or an alarm cue were delivered to the demonstrator in dyadic tests involving familiar and unfamiliar pairs.

### Distance to the conspecific’s tank (proximity to the demonstrator)

Following the addition of the alarm substance to the demonstrator, the observer swam closer to the demonstrator’s tank (**Figure 5**, estimate = 0.60, *P* < 0.001), something that did not happen when distilled water was added. Distance to the demonstrator decreased over basal values (estimate = 0.43, *P* = 0.005) but increased with familiarity (estimate = 0.16, *P* = 0.031) and was also affected by the interactions between demonstrator and time (estimate = 0.36, *P* = 0.03), and between time and stressor (estimate = −0.49, *P* < 0.001). There was thus evidence of fear contagion with respect to proximity to the other fish’s tank and this also seemed to be influenced by familiarity.

**Figure 5.**
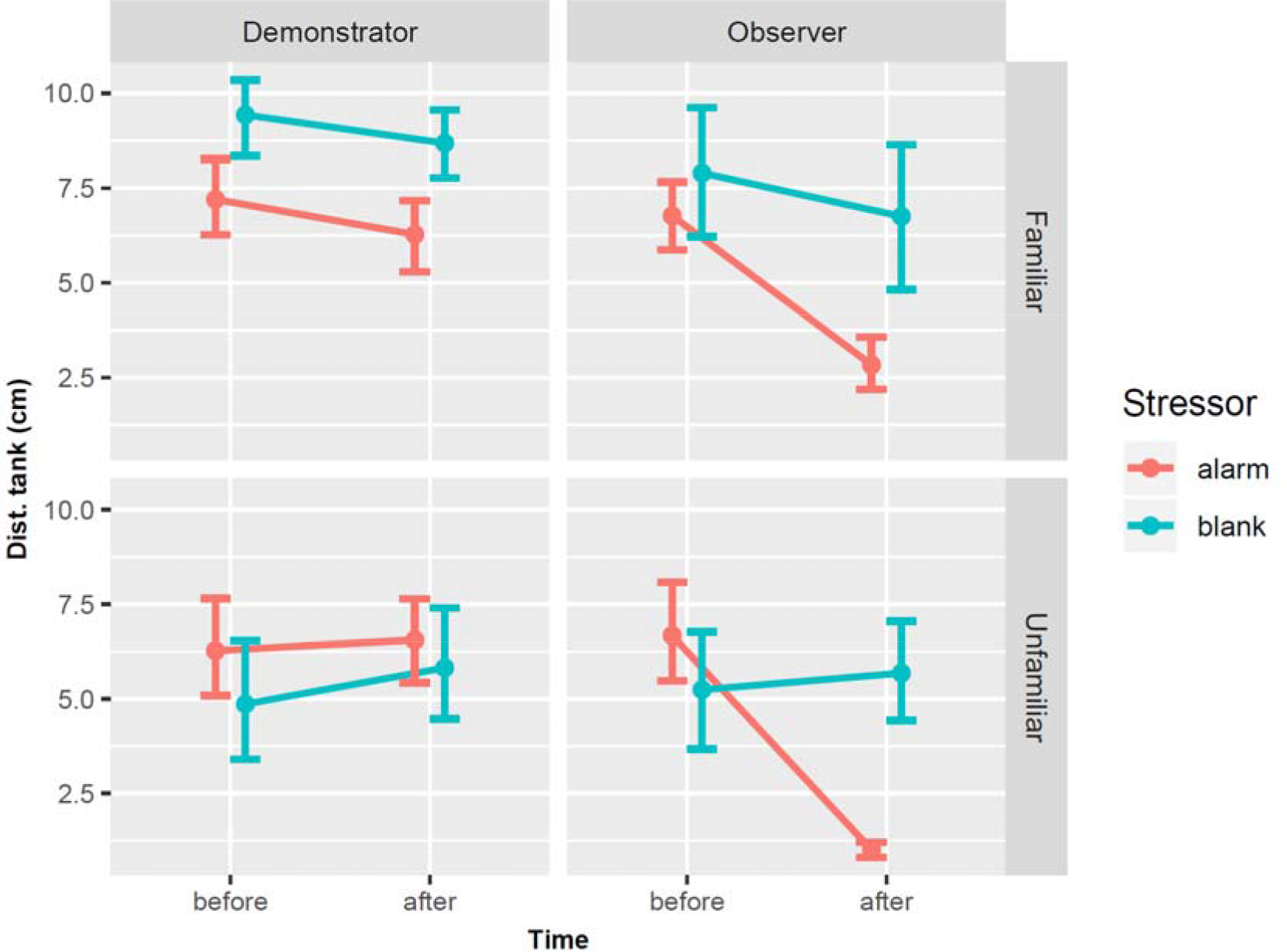
Changes in the distance to the conspecific’s tank (mean ± 95CI) of zebrafish before (basal) and after distilled water or an alarm cue were delivered to the demonstrator in dyadic tests involving familiar and unfamiliar pairs.

### PCA analysis

PCA analysis showed that the behaviour of demonstrators and observers was similar and showed little variation among individuals when the fish were not stressed (i.e. when distilled water was added or before an alarm cue was added, **Figure 6a-c**, **Figure 6e-g**). The first component, PC1, accounted for 89.8% of the variation and described freezing behaviour, while PC2 accounted for 5.9% of the variation and described variation in swimming speed and proximity to the conspecific. No statistical difference between observer and demonstrators was found along PC1 (*F*_3,90_ = 0.847, *P* = 0.472), but there was a marked difference along PC2 (*F*_3,90_ = 68.52, *P* <0.001; **Figure 6d, Figure 6h**) which depended on the type of fish (estimate for observer = 1.21, SE = 0.64, *P* = 0.009), the extent of familiarity (estimate for unfamiliar = −7.72, SE = 0.91, *P* < 0.001), and their interaction (estimate for observer x unfamiliar = 15.44, SE = 1.29, *P* < 0.001). Thus, observers were better able to mimic the anti-predatory behaviour of demonstrators when they were familiar than when they were unfamiliar (**Figure 6d)**.

**Figure 6.**
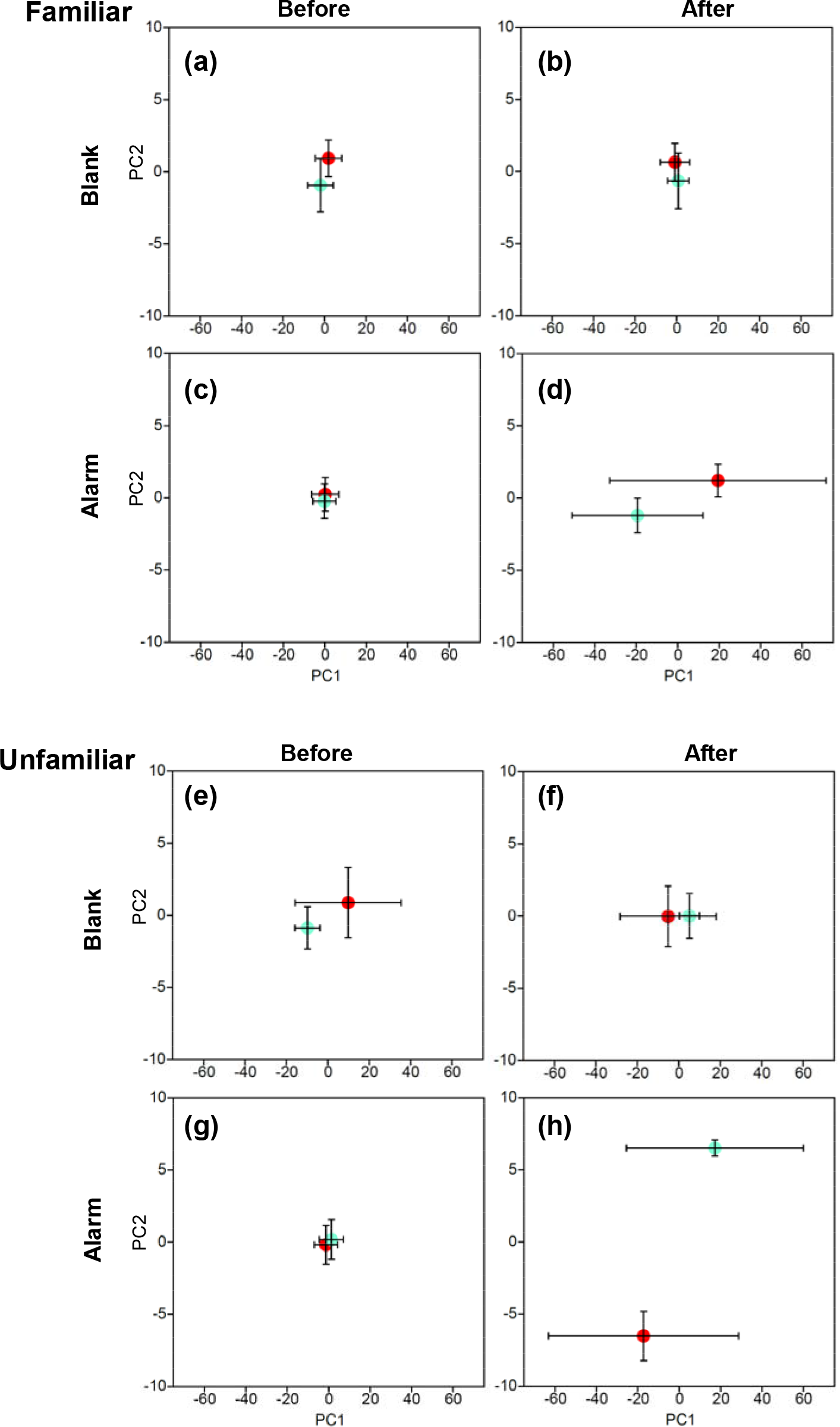
Variation along the first two principal components (means ± 95CI) describing the behaviour of familiar and unfamiliar dyads of zebrafish before (basal) and after distilled water or an alarm cue were delivered to the demonstrator (red) but not to the observer (green).

## DISCUSSION

Our study provides novel evidence in support for the existence of fear contagion in fish, and suggests that this is influenced by familiarity, as shown for mammals (Gonzalez-Liencres, et al 2014; Jeon et al., 2010). The alarm response of zebrafish has been characterised by an increase in swimming speed interspersed with freezing bouts and diving to the bottom (Kalueff et al., 2013). To trigger a fear response we added an alarm substance to the water, *schreckstoff*, (von Frisch, 1938), a well-established stressor for zebrafish (Jesuthasan & Mathuru, 2008). As expected, the addition of the alarm substance triggered a flight response in zebrafish, which swam faster, moved to deeper waters, and included bouts of freezing behaviour, something not observed when distilled water was added. Such anti-predatory behaviours were mimicked to some extent by the observers, even though they had no direct exposure to the stressor. Thus, the mere sight of a distressed conspecific was enough to trigger in the observer the same fear response experienced and displayed by the demonstrator.

Our study shows that fear contagion is modulated by familiarity in zebrafish, since three of the four responses examined varied depending on whether the observers were watching individuals they were familiar with. Compared to unfamiliar demonstrators, observers responded to the sight of familiar demonstrators by matching their swimming speeds more closely, and by moving closer to the bottom. Observers also moved closer to the demonstrator at the sight of a distressed conspecific, but - perhaps unexpectedly, this was more pronounced when they were paired with unfamiliar demonstrators. These results add support to evidence indicating that fish are capable of identifying and reacting to the behaviour of conspecifics (Jesuthasan & Mathuru, 2008; Rey et al., 2015). In social species, fear contagion is thought to be adaptive as it will often increase the probability of escaping from predators (Nakahashi & Ohtsuki, 2018).

Among fishes, several studies have shown that familiarity increases shoal cohesion (Chivers et al. 1995; Lachlan et al. 1998) and facilitates social learning (Swaney et al., 2001), and our study shows that familiarity also affects fear contagion, which may explain why association with familiar fish is generally adaptive (Griffiths, 2003). For example, familiar brown trout respond 14% faster than unfamiliar fish to a predator attack (Griffiths et al. 2004), most likely because they can interpret signals from familiar fish more accurately. Familiarity in our study was established rapidly, after only seven days of cohabitation, which is consistent with previous results in zebrafish (Madeira & Oliveira, 2017) and other species (Griffiths, 2003), where individuals were able to recognize familiar neighbours after short periods of interaction. Although it is possible that some individuals in our study may have become familiar in the stock tanks (i.e., before the 7 day cohabitation experiment), this is unlikely as the group size was too large (100 fish/tank) for individual recognition (Griffiths & Magurran, 1997), fish were mixed and allocated at random to 26 groups, and this would not explain why familiar and unfamiliar fish behaved so differently.

The advantages of familiarity may be accrued through visual recognition, but also through chemical cues (Griffiths, 2003), as fish can recognize the metabolites of conspecifics released in the water (Ward et al. 2009). Zebrafish can use both chemical and visual cues for individual recognition (Hinz et al., 2013), but our study indicates that visual signals alone are enough to trigger fear contagion, as familiar fish behaved differently from unfamiliar fish when no chemical signals were exchanged between observers and demonstrators. The strong shoaling behaviour of zebrafish may help explain the evolution of fear contagion on this species. Unlike social learning, which is thought to have evolved to facilitate the long-term transmission and storage of information (Brown & Laland, 2003), fear contagion may have evolved to deal with rapid, short-term signals and swift responses, such as the anti-predatory response (Nakahashi & Ohtsuki, 2015).

It has been proposed that behavioural contagion should be heightened when the demonstrator displays abnormal or extreme behaviours (Nakahashi & Ohtsuki, 2015). Freezing is an extreme behaviour that can be induced by alarm substances and is commonly seen in many fish species in response to predators (Gerlai et al. 2000; Gerlai & Csányi 1990; Miklosi el al. 1997; Roberts et al. 2011; Roberts & Garcia de Leaniz, 2011). In this sense, the increased duration in freezing bouts following exposure to the alarm substance was highly contagious in our study, but it was not affected by familiarity. We suggest that for zebrafish, freezing behaviour constitutes a more robust signal of danger than bottom dwelling or an increase in swimming speed. Hence, it may be adaptive for an individual to freeze when another one is freezing, regardless of the sender’s identity. On the other hand, changes in swimming speed or in the position in the water column form part of the normal behaviour of zebrafish (Kalueff et al., 2013) and may represent less extreme, and hence less reliable, signals.

Our results show that zebrafish can not only distinguish between familiar and unfamiliar conspecifics by visual cues alone, but that they can also eavesdrop on their neighbours living in separate tanks and adjust their behaviour accordingly. This may have implications for fish welfare if, for example, fish can become stressed simply by watching their neighbours become stressed. In the wild, eavesdropping may be adaptive as it allows zebrafish to acquire information on predatory threat from shoal neighbours (Abril-De-Abreu et al., 2015; Oliveira et al., 2017), but the implications for fish welfare in captivity deserve further attention. In livestock, contagion of negative emotions such as fear and anxiety can impair the behaviour and health of the group (Reimert et al., 2013), and our study suggests that the same could happen in zebrafish.

Our results also have implications for experimental design because fish housed in separate tanks may not be assumed to be statistically independent (Colegrave & Ruxton, 2017) if their behaviour is affected by that of others. Visual isolation of tanks, therefore, must be ensured to prevent eavesdropping. Ultimately, our study indicates that fish are capable of recognizing and mimicking individual behaviours and place them into the right context. Future studies might benefit from investigating if behavioural contagion also occurs in relation to positive stimuli (such as access to food, mates, or enriched habitats) as this could perhaps be used to improve welfare.

Declaration of interests: none

## ACKNOWLEDGEMENTS

We are grateful to Maria E Leite and Julia Ruiz-Oliveira for assisting with the collection of data. This study was funded by a Royal Society Newton Mobility grant to Prof Sonia Consuegra, ACL, PFS and CGL and the ERDF SMARTAQUA Operation.

