## Supplementary material for "Fear contagion in zebrafish: a behaviour affected by familiarity": Supp Fig S1

**Figure S1.** Temporal variation in the response of single zebrafish (i.e. demonstrators without observers) to the addition of either distilled water (blank) or an alarm cue . Shown are means ± 95 CI.

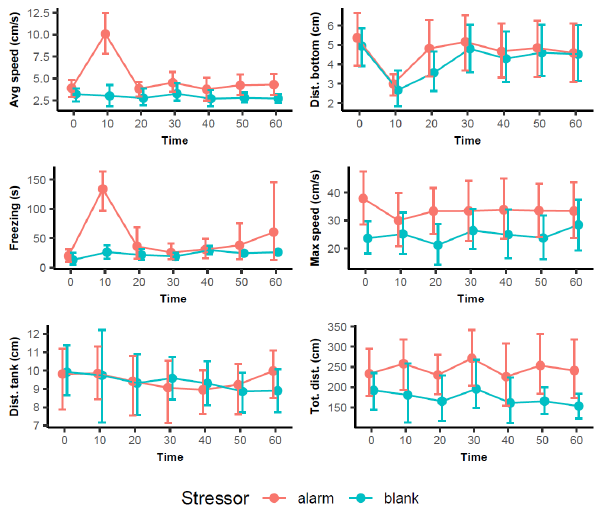
