## Supplementary material for "Fear contagion in zebrafish: a behaviour affected by familiarity": Supp Table S1

**Table S1.** Behaviour of familiar zebrafish held with and without a partition during the acclimatization period. Values represent means ±SD.

|  | **Demonstrator** | | | | | | **Observer** | | | | | | |
| --- | --- | --- | --- | --- | --- | --- | --- | --- | --- | --- | --- | --- | --- |
|  | Basal | |  | 10 min | |  | Basal | |  | 10 min | |  | |
| Behavioural parameter | no cover | cover | *p* | no cover | cover | *p* | no cover | cover | p | no cover | cover | | *p* |
| Average speed (cm/s) | 2.6 ± 0.9 | 3.3 ± 2.3 | 0.07 | 4.7 ± 3.0 | 4.1 ± 1.6 | 0.82 | 3.0 ± 1.3 | 3.1 ± 1.9 | 0.26 | 6.2 ± 1.7 | 3.8 ± 5.1 | | 0.34 |
| Maximum speed (cm/s) | 42.1 ± 14.8 | 53.7 ± 12.5 | 0.05 | 39.2 ± 27.9 | 44.0 ± 16.4 | 0.66 | 41.6 ± 14.9 | 46.5 ± 13.3 | 0.44 | 28.1 ± 18.5 | 32.8 ± 15.8 | | 0.53 |
| Total dist. travelled (cm) | 199.3 ± 110.3 | 294.1 ± 100.9 | 0.07 | 192.7 ± 107.5 | 250.1 ± 75.8 | 0.23 | 199.5 ± 134.1 | 263.2 ± 105.7 | 0.26 | 161.9 ± 116.2 | 190.4 ± 56.7 | | 0.54 |
| Freezing (s) | 27.9 ± 16.8 | 15.5 ± 9.9 | 0.08 | 192.4 ± 136.9 | 180.5 ± 38.9 | 0.82 | 28.2 ± 13.9 | 16.9 ± 9.1 | 0.05 | 153.6 ± 81.5 | 154.7 ± 22.7 | | 0.97 |
| Distance from bottom (cm) | 5.1 ± 2.9 | 3.9 ± 2.8 | 0.34 | 2.0 ± 1.8 | 1.4 ± 0.4 | 0.38 | 5.5 ± 3.1 | 3.1 ± 1.7 | 0.06 | 1.4 ± 1.0 | 1.9 ± 0.6 | | 0.19 |
| Distance to other tank (cm) | 7.2 ± 2.6 | 5.5 ± 0.9 | 0.11 | 6.3 ± 2.5 | 6.6 ± 2.9 | 0.79 | 6.8 ± 2.3 | 6.3 ± 1.8 | 0.64 | 2.8 ± 1.7 | 3.0 ± 2.3 | | 0.83 |

p-values corresponds to the student t tests between groups held with or without cover.
